# High-resolution 4D spatiotemporal analysis reveals the contributions of local growth dynamics to contrasting maize root architectures

**DOI:** 10.1101/381046

**Authors:** Ni Jiang, Eric Floro, Adam L. Bray, Benjamin Laws, Keith E. Duncan, Christopher N. Topp

## Abstract

Root systems are branched networks that develop from simple growth properties of their individual roots. Yet a mature maize root system has many thousands of roots that each interact with soil structures, water and nutrient patches, and microbial ecologies in the micro-environments surrounding each root tip. Although the plasticity of root growth to these and other environmental factors is well known, how the many local processes contribute over time to global features of root system architecture is hardly understood. We employ an automated 3D root imaging pipeline to capture the growth of maize roots every four hours throughout seven days of seedling development. We model the contrasting architectures of two maize inbred genotypes and their hybrid to derive key parameters that distinguish complex growth patterns as a function of time. The statistical characteristics of local root growth defined the global system properties despite a large range of trait values. “Computational dissection” of a single root from each root system identified differences in the size of the root branching zone and lateral branching densities, but not radial patterns, that drove the contrasting root architectures from seedling to maturity. X-ray imaging of mature field-grown root crowns showed that seedling growth trajectories persisted throughout development and could predict eventual architectures, suggesting a strong genetic basis. The work connects individual and systemwide scales of root growth dynamics, providing the means for a function-valued approach to understanding the genetic and genetic x environment conditioning of root growth that will enable breeding for enhanced root traits.

**SIGNIFICANCE STATEMENT:** When and where roots grow determines their ability to capture short-lived and patchy water and nutrient resources to support the aboveground organs of the plant. Roots have no known long-distance external sensing mechanisms, but form branched networks that blindly explore the soil and respond to encountered local stimuli. How global architectures form from the many thousands of these local responses, and how they are controlled genetically are major open questions. Here we quantify differences in local root growth patterns of two inbred genotypes of maize that control contrasting systemwide properties. Measurements at the seedling stage were highly correlated with the complex architectures of mature root systems, paving the way for the development of crops with greater resource uptake capacity.

## INTRODUCTION

Root system architecture (RSA) is the foundation of plant growth and productivity (1). It affects water and nutrient acquisition efficiency (2, 3), and can change in response to different environments (4, 5). Understanding the genes and mechanisms that govern root system architecture would be a major new lever for crop improvement work (6–8). However, while genetic studies of shoot growth have benefited from automated and dynamic quantification (9, 10), the spatial and temporal aspects of root system architecture, and their genetic basis, remain obscure.

Root systems begin with a single primary root. In many monocots, and especially the annual grasses such as maize, additional seminal roots can subsequently emerge from the seed (11). Later, nodal roots emerge in whorls from shoot derived tissue. All of these root types grow exponentially in size and complexity from three simple processes local to the root tip: elongation, curvature, and lateral branching. A 3D network forms that in maize can ultimately consist of tens of thousands of roots occupying over 200 cubic feet of soil (11). These global architectural properties of the root system determine the ability of the plant to capture edaphic resources in the heterogeneous and dynamic environments characteristic of soil.

Intensive research has yielded detailed molecular and cellular mechanisms of how single roots grow at the local scale (reviewed in: (12, 13)) on one hand, and the identification of global architectures, or ideotypes, that are best suited for resource capture in natural or agricultural environments (14–17) on the other, but rarely have the two been experimentally connected. Structural-functional models can simulate a range of root architectures based on equations programmed to reproduce local growth and environmental interactions (18–20), and have been used to predict empirical data (21, 22). However realistic parameterization and constraint of these models is thus far piecemeal, lacking adequate empirical data sets that incorporate time dynamics and genetically encoded differences in root development and genetic x environment interactions. A phenotyping system that bridged both local and global root growth dynamics would capture the process of how root architecture complexity builds through many iterative and partially stochastic local patterns.

Image-based phenotyping technologies that enable non-destructive measurement of root traits have been developing at a rapid pace in recent years (23, 24). Minirhizotron tubes have been used to measure roots in the field at multiple time intervals (25), but have the limitation of only examining the roots growing along the transparent tube. GROWSCREEN-Rhizo uses soil-filled rhizotrons to image visible roots though transparent 2D plates (26). Time-lapse 2D imaging allow the dynamics of root growth and environmental responses to be studied (27–35), but these studies are dimensionally constrained. To analyze whole root systems in 3D, platforms where roots grow in gel cylinders have been developed (36–39). X-ray computed tomography (40, 41) and magnetic resonance imaging (42–44) also provide 3D quantification of roots in soil. Manually repeating 3D acquisitions at different time points enables the study of unconstrained root growth (36, 38, 45–47), but at long time intervals and with only global quantifications of root architecture. Although at least one 3D imaging approach has been used to identify SNPs that corresponded to yield increases in field experiments (48), it has been pointed out that few genes identified by lab-based reverse genetics have been corroborated in the field (49), causing some to call into question the usefulness of non-field-based root work for applied purposes (50).

In this paper, we have developed an optical imaging system that allows frequent 3D monitoring of root growth. Using automatic time lapse imaging, growing root systems of B73, Mo17, and their hybrid were monitored over one week of development until day 11 after germination (DAG). Using computer vision, we computed the time function for each root and analyzed the growth patterns. Mathematical modelling of the time series revealed the key parameters that drive genotype-specific differences in root architecture, including timing of a sharp inflection point in relative growth rate. 3D analysis of field grown-roots showed these patterns persist to maturity and are thus more influenced by genetics than the environment. Our high-resolution spatiotemporal imaging and analysis approach will facilitate the study of growth responses to resource patches, other roots, soil particles, or other heterogeneously distributed soil parameters at both the local and global level. This work will enhance the development of empirically-driven, probability-based growth models that can accurately predict root growth and root-environment interactions as a function of genotype.

## RESULTS

Plant roots have typically been studied either individually (locally) in great detail, or globally as entire systems, but rarely both, especially as freely-growing 3D structures. In order to understand how local root growth patterns contribute to global architectures along both a real-time and a developmental time axis, we established a 4D analysis pipeline (Figure 1). We automated a 3D gel-based optical imaging system (39, 45) for time lapse, and paired it with DynamicRoots software (46), and custom R code (available on Github: https://github.com/Topp-Roots-Lab/timeseries_analysis) to get the dynamic traits of individual branches based on the time series of 3D shapes. Two historically agronomically important maize inbred genotypes with contrasting root architectures, B73 (a stiff-stalk) and Mo17 (a non-stiff-stalk), and their hybrid were imaged every 4 hours across 8 days of development (164 hours total), from day 4 to day 11 after germination. The plants typically had only a primary root and one or two seminal roots at the beginning of the experiments, but over one hundred root branches by the end, including nodal roots (Movie S1), providing a wide spectrum of architectural complexity.

**Figure. 1.**
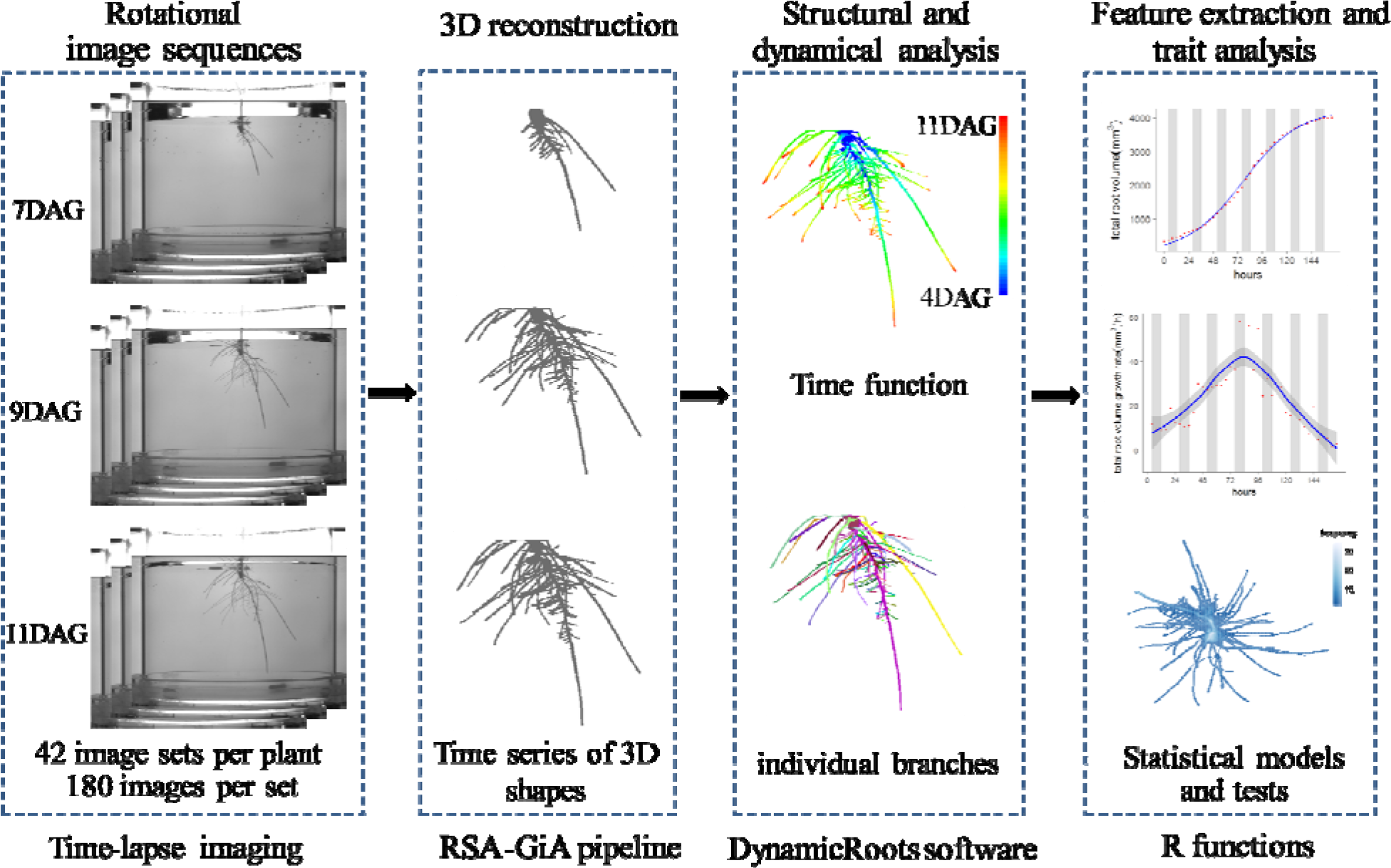
Workflow for 4D analysis of maize roots. Rotational image sequences from our automatic 3D time lapse imaging system were imported and processed to reconstruct 3D models using the RSA-GiA pipeline. The reconstructed time series of 3D models were processed using DynamicRoots software. The phenotypic features were extracted and analyzed using custom R code.

### Quantifying seedling global root traits at fine temporal scales reveals fundamental differences in growth patterns between two maize inbred genotypes and their hybrid that persist through maturity

Phenotypic analysis of root architecture in a panel of diverse maize inbred lines has shown complex genotype-specific patterns that change during early development (39), but their expression over time and the underlying cause is not yet understood. Using the automated imaging system, we were able to observe global-scale growth patterns on a highly resolved temporal scale. To provide information about the size and shape of the root systems, we compared total root volumes, lengths, and numbers, which all increase over time (Figure 2). Despite a highly controlled and homogeneous environment, the dynamic range of growth varied extensively among individual plant as evidenced by the large spread of data. Yet phenotypic separation of the two genotypes could be identified very early in the experiment along the values of the mean curve (Figure 2a-c; t-tests for each time point in Table S1-S3). The hybrid values had the greatest intragenic variation, but the average values were largely intermediate, providing no evidence for heterosis in seedling root growth, similar to previous findings (51). While the total root volume and number of roots in B73 samples were consistently greater than Mo17 at all time points, initial differences in total root length disappeared by day 11 (Figure 2, Table S2). Since dry root weights from day 11 also showed that B73, Mo17 and their hybrid all had similar root biomass (Figure S1), we conclude that Mo17 and B73 fundamentally differ in how they allocate carbon resources for root foraging. B73 invests in relatively more, shorter, and “cheaper” roots (i.e. less biomass per unit of surface area), and Mo17 invests relatively more in the continued growth of extant roots, with their hybrid as intermediary.

**Figure 2.**
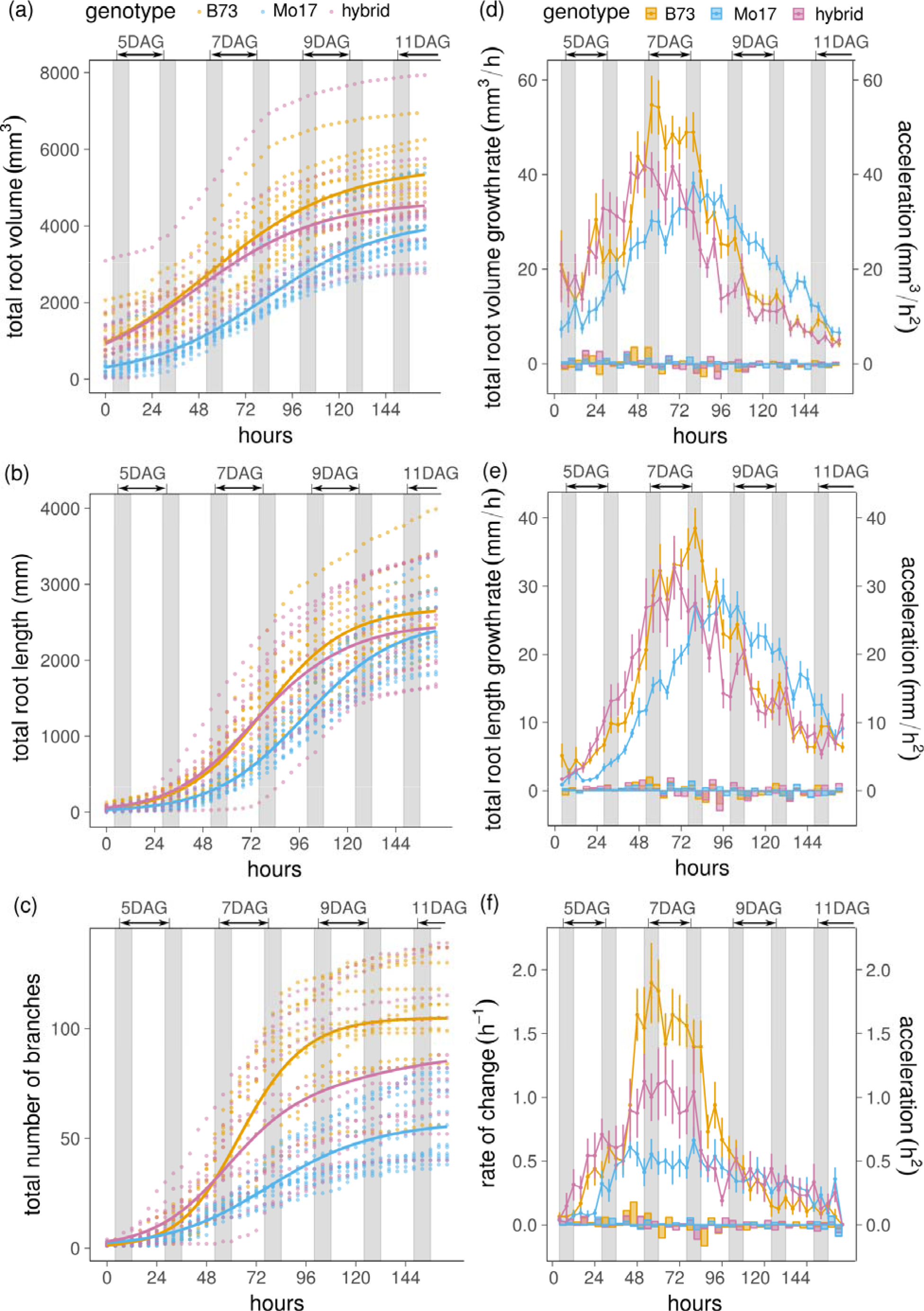
Global root trait dynamics reveal fundamental differences in growth patterns. Timecourse values of total root volume (a), total root length (b), and total root number (c). Each point represents an individual seedling, and the solid lines indicate mean values. Timecourse of growth rates and accelerations of the total root volume (d), total root length (e), and total root number (f). Points represent the mean value of growth rates. Vertical error bars represent the standard errors. Barplots show the accelerations. The gray bars represent night time.

To capture the underlying dynamics of these temporal relationships, we computed the rates of change (or velocities, the first derivative) and accelerations (changes in velocity, the second derivative) for the three global traits (Figure 2d-f). A sharp demarcation occurs in both inbred genotypes that separates increasing and decreasing growth rates for root volume and length. However the inflection point is delayed in Mo17 relative to B73 (Figure 2d,e), and slightly earlier in the hybrid. This transition is coupled to the rate of new roots in B73 and the hybrid, but not in Mo17 (Figure 2f), supporting evidence for a fundamentally different process for patterning root architecture between the two genotypes that appears semi-dominant in the hybrid.

Despite the clear trend over the 1-week time interval, accelerations or decelerations between any two 4-hour time intervals were generally smooth - no significant differences were found for rates of change in total root number for any genotype at this temporal resolution, and in only a single interval each in B73 and the hybrid for root volume. Rates of change for total root length were more variable in all genotypes, but not in a consistent pattern (Figures 2d-f; Table S4). Diurnal patterns of growth in plant leaves and shoots (52) are well established, but evidence in grasses suggests that their roots do not change elongation patterns along day/ night cycles (27, 28). We compared the growth rates during the 4 hours before dark to the growth rate during the next 4 hours in the dark for each genotype. All three genotypes had differences (*p* < 0.05) in day/ night growth rates at some time points, but not others (Table S4), providing no strong evidence of diurnal regulation, which reinforces previous studies. We found that growth rates were predominantly driven by developmental time, rather than diurnal real-time.

We investigated the duration and environmental conditioning of these apparent genotype-specific properties by extracting identical 3D features from X-ray scanned root crown samples (53) excavated from a field at anthesis (Figure 3). The phenotypic trends of the inbreds are remarkably consistent, with nearly every significant difference between B73 and Mo17 in the gel-grown seedlings also reflected in the field-grown mature root systems (Figure S2), including total root volumes, lengths, and numbers. In multivariate space, these high-resolution 3D phenotypes clearly delineated the two inbred genotypes along similar eigenvector distributions, with the first two principal components (PCs) explaining ~80% of the total phenotypic variation in each data set (Figure 3c, d; Table S5). As expected, the aboveground biomass of field-grown hybrids was much larger than either parent (Figure S1), but the expression of heterosis was detected to a lesser extent in the 3D root data, with extreme values for only total root volume and convex hull area. Nonetheless, the first two PCs separated the hybrid from either parent in the field data (Figure 3d). Contrary to conventional wisdom, we find a strong correspondence between maize seedling and mature root traits. Our results suggest that, given sufficiently high-quality data, a strong and persistent genetic influence on root growth patterns can be identified across time and environment.

**Figure 3.**
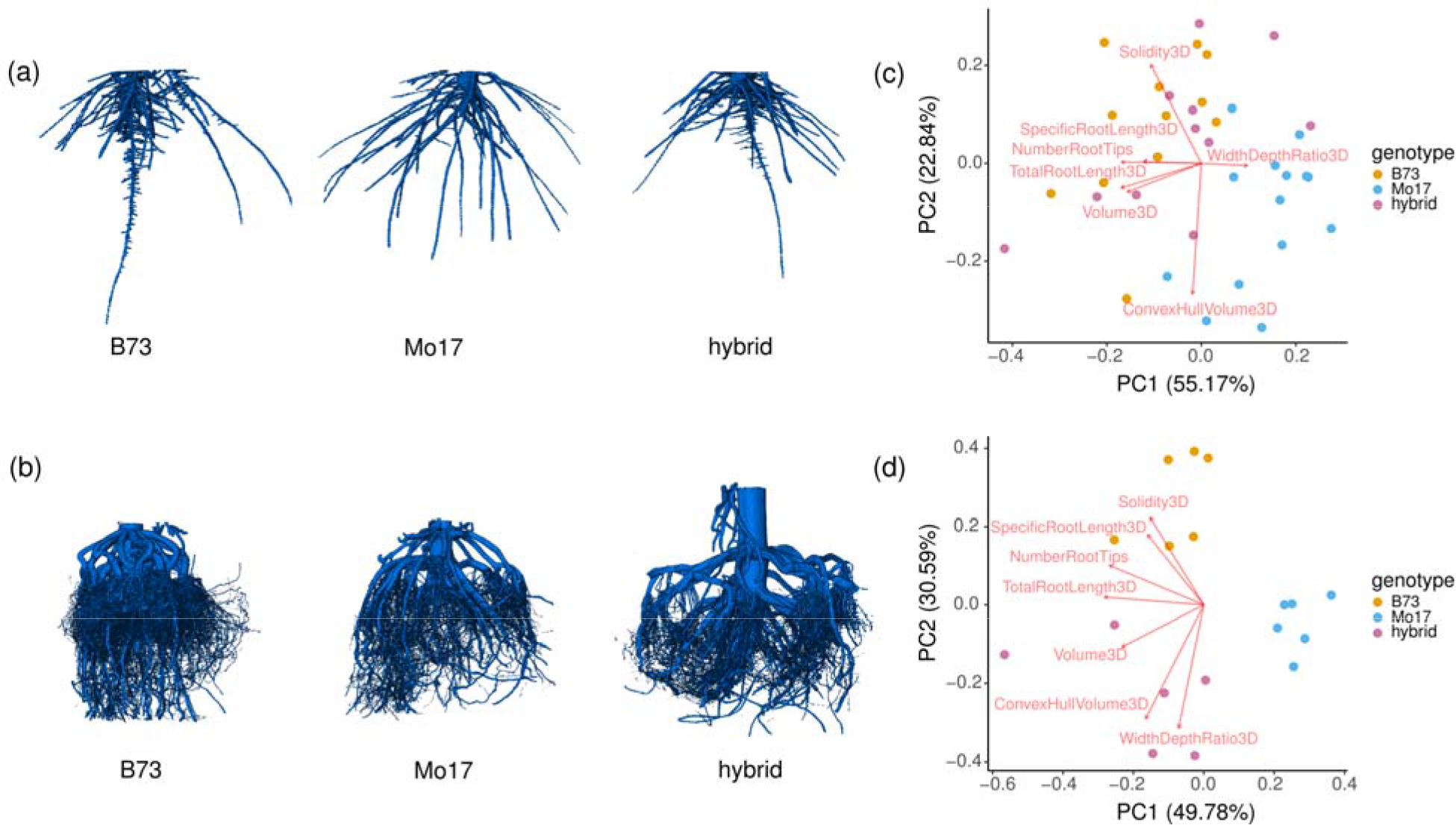
Differences in seedling root systems persisted in mature root systems. 3D reconstructions of seedling root systems at 11DAG (a), and mature root crowns at anthesis (b). Score and loading plots for the first two principal components (PC) of seedling root traits (c), and mature root crown traits (d). Each dot represents one plant.

### Modelling global growth patterns allows a direct comparison of key parameters that control more complex patterns

Non-linear models such as logistic growth functions can provide “function-valued” traits that integrate both real and developmental time to describe complex patterns of growth or architectural change (54–57). After testing various nonlinear models for several fitting criteria (Table S6), we chose a 3-parameter logistic growth function to model the global trait dynamics (Figure 4; Methods). Parameter α_1_ describes the “carrying capacity” of the function, which reports the maximum values of the modelled growth curve. B73 had significantly higher α_1_ values for total root volume and total number of root branches, reflecting that B73 has a larger final volume and more branches than Mo17 (Figure 4). However, the α_1_ values for total root length were not different, despite significantly larger values for B73 at each time point until hour 144. This result highlights the importance of capturing and integrating the time dimension through an appropriate model, rather than ad hoc analysis of individual time points, which may be misleading. Parameter α_2_ represents the maximum rate of change and is a direct measure of the trends seen in Figures 2 d-f. It provides an explicit quantification of the different times at which the inflection points between increasing and decreasing growth trends were reached for a given trait. B73 and the hybrid begin slowing the rate at which root length and volume are added more than 24 hours prior to Mo17 (Figure 4a,b), whereas differences in the time at which additions of new lateral roots slowed were much less pronounced (Figure 4c). Parameter α_3_ captures the slope of the growth curve, and except for total root volume, the values between B73 and Mo17 were significantly different, reflecting that the total number of branches and to a lesser extent total root length increased more sharply for B73. The overall growth trends for the hybrid were most similar to B73, although we note that range of values was typically larger than either parent (reflected in the long axes of the violin plots, Figure 4), suggesting the hybrid is less constrained in its developmental program.

**Figure 4.**
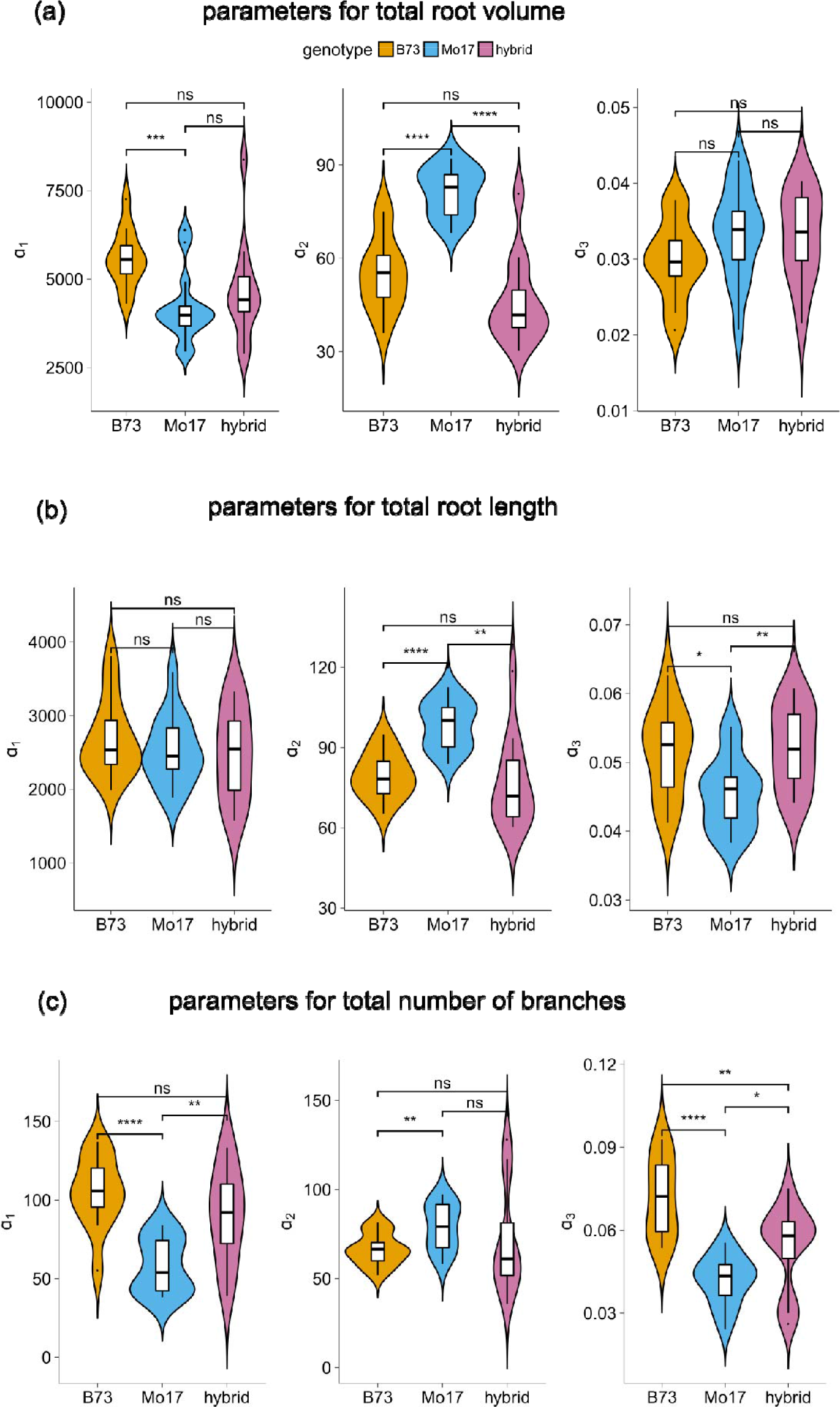
Comparison of global growth patterns by modelling. Parameters (α_1_, α_2_, α_3_) of the logistic growth models estimated for total root volume (a), total root length (b), and total root number (c). Violin plots and box plots were generated using ggplot2 package with default settings for statistics in R. The significance levels of pairwise t tests were added. (ns: p>0.05, *: p<=0.05, **: p<=0.01, ***: p<=0.001, ****: p<=0.0001)

Modelling of dense (4-hour intervals) time-series data provided important insights into the growth dynamics underlying genotypic differences in root architecture, but from a practical standpoint, we wished to know if similar answers could be derived from fewer data points. We conducted two types of sensitivity tests for each trait by progressively removing data from the analysis to generate pseudo time intervals and either: 1. Comparing the differences in modelled trait values for a single genotype (B73, Figure S3-5), or 2. Comparing the differences in statistical test results among B73, Mo17, and the hybrid (Figure S6-8). Our estimates of the B73 total root volume modelling parameters would not have changed significantly if we would have imaged at any interval of 20 hours or less, whereas 4 versus 24-hour intervals gave statistically different results. Model estimates of total branch number hardly varied at pseudo time interval from 4 to 24 hours with a few sporadic exceptions. However, total root length estimates were extremely labile, resulting in significant differences in the values between 4 hours and nearly every other interval (Figure S3-5). Despite this intra-genotypic variability, the computed statistical differences among B73, Mo17, and the hybrid were remarkably consistent at any time interval from 4 to 24 hours for all three traits (Figure S6-8). The results of this analysis suggest that much of the same information about root growth dynamics could be attained through less frequent phenotyping, which could translate to increases in sample throughput, and more power to resolve subtle environmental or genetic differences.

### Analysis of the time function reveals local growth patterns driving genotypic differences in root architecture

Root architecture is a global property determined by the cumulative elongation, branching, and curving of each root. Yet much is unknown as to how these local parameters emerge as global root system properties, especially in the context of three-dimensional complexity. A key feature of our phenotyping pipeline is automated derivation of the time function and quantification of local growth (Methods). We analyzed all of the individual branches as aggregate trait distributions during the timecourse, from which fundamental differences in growth patterns between genotypes were discerned (Figure 5). Analysis of root length distributions showed that while B73 had a higher proportion of longer branches than Mo17 at the earliest time points, the proportion of shorter branches increased rapidly for B73 during the 5-7 DAG (Figure 5a). This trend reflects the rapid proliferation of lateral roots captured in the global architecture and growth modelling analyses (Figure 2,c,f and 3c). However the burst of new roots quickly tapered by 8DAG as B73 increasingly elongated existing roots, rather than forming new ones. This fundamental shift in resource allocation is clearly seen by the steadily decreasing proportion of relatively shorter roots throughout the rest of the experiment (Figure 5a). In comparison, the architecture of Mo17 was initially defined by shorter roots, but then consistently balanced existing root elongation and new root production throughout the timecourse, reflected in the steady root length distributions seen from 8DAG. Growth patterns of the hybrid were the most consistent, but there was a greater emphasis on new root production relative to Mo17.

**Figure 5.**
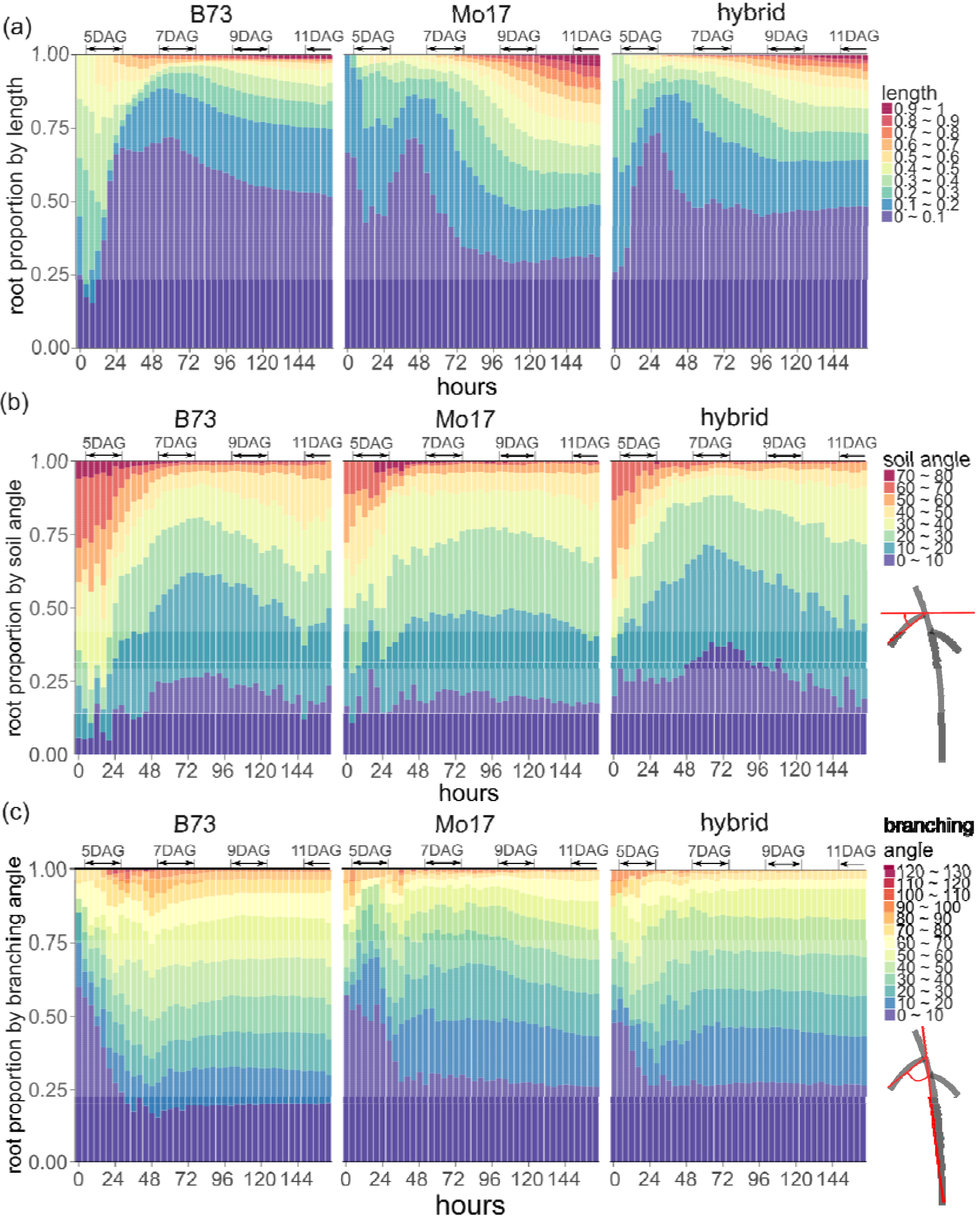
Local growth patterns revealing genotypic differences. (a) Timecourse of root length distribution. Root length for each individual branch was normalized by the longest branch in the whole root system and recorded in 10 bins. (b) Timecourse of soil angle distribution. The soil angle was the angle formed by the root branch and the horizontal level. (c) Timecourse of branching angle distribution. The branching angle was the angle formed by the child branch and its parent branch.

To study root curving, we computed the soil angle distribution, which captures root geometry relative to an extrinsic reference (the soil horizon), and branching angle distribution, which measures the intrinsic angle between a child and parent branch regardless of orientation to the soil horizon. As shown in Figure 5b, B73 had relatively larger soil angles relative to Mo17 at the early time points, indicating that B73 root systems begin growing at a steeper angle. However the proportion of shallow-angled branches quickly increased with the emergence of new branches (Figure 2c), peaking at 7DAG for B73 and the hybrid, but Mo17 was quite consistent during the experiment. Since most roots eventually orient to the vertical gravity vector over time, the production of new lateral roots and their angles relative to the parent branch (which is older and thus more likely to be vertically-oriented) largely dictate the extent of horizontal soil exploration. In the earliest stages of maize root development, several embryonic (seminal) roots emerge from the seed, which is reflected in all three genotypes by the dominance of shallow root branching angles at the earliest time points. As the root system architecture became more complex over time, branching angle distribution patterns were consistent, but the magnitudes depended on genotype (Figure 5c). The relatively greater angles for B73 demonstrate that lateral roots grew further away from the parent branch than for Mo17 and the hybrid, highlighting fundamental differences in local growth patterns that have strong implications for root architecture and resource capture (58).

### High-resolution analysis along a single root reveals developmental processes driving genotypic differences in root architecture

Genetic variation for intrinsic developmental processes results in an astonishing diversity of plant morphology. Single roots have been used extensively to study plant development, in part because the position of lateral roots are arranged longitudinally across time, with the youngest organ always nearest to the growing tip at the beginning of the maturation zone (59). To understand how genotypic differences in root development contribute to the emergence of complex architectures, we computed the branch hierarchies for entire root systems, enabling us to know the precise topological relationships of all the roots in each 3D model (Methods). For the final, most architecturally complex time point, we defined the primary root as the branch connected to the seed which had the most child branches, and analyzed the patterns along this developmental axis. The positions of each lateral root were computed using the distance between the branching fork and the base of the primary root, divided by the primary root length (Figure 6a). B73 had more total lateral branches along the primary root than Mo17, but they also emerged much closer to the root tip. On average, B73 lateral roots were found along the upper 80% of the primary root, whereas Mo17 lateral roots were found along only the upper 60%. Since there was no significant difference in the primary root length between the two genotypes, B73 had both a higher lateral root branching density and a longer zone of branching. These data are parsimonious with the local and global root length and branching traits we measured previously (Figures 2 and 5). Thus, the basis for the architectural tradeoff of investment in new branches versus growth of existing branches is apparently controlled by genetic differences in local developmental processes.

**Figure 6.**
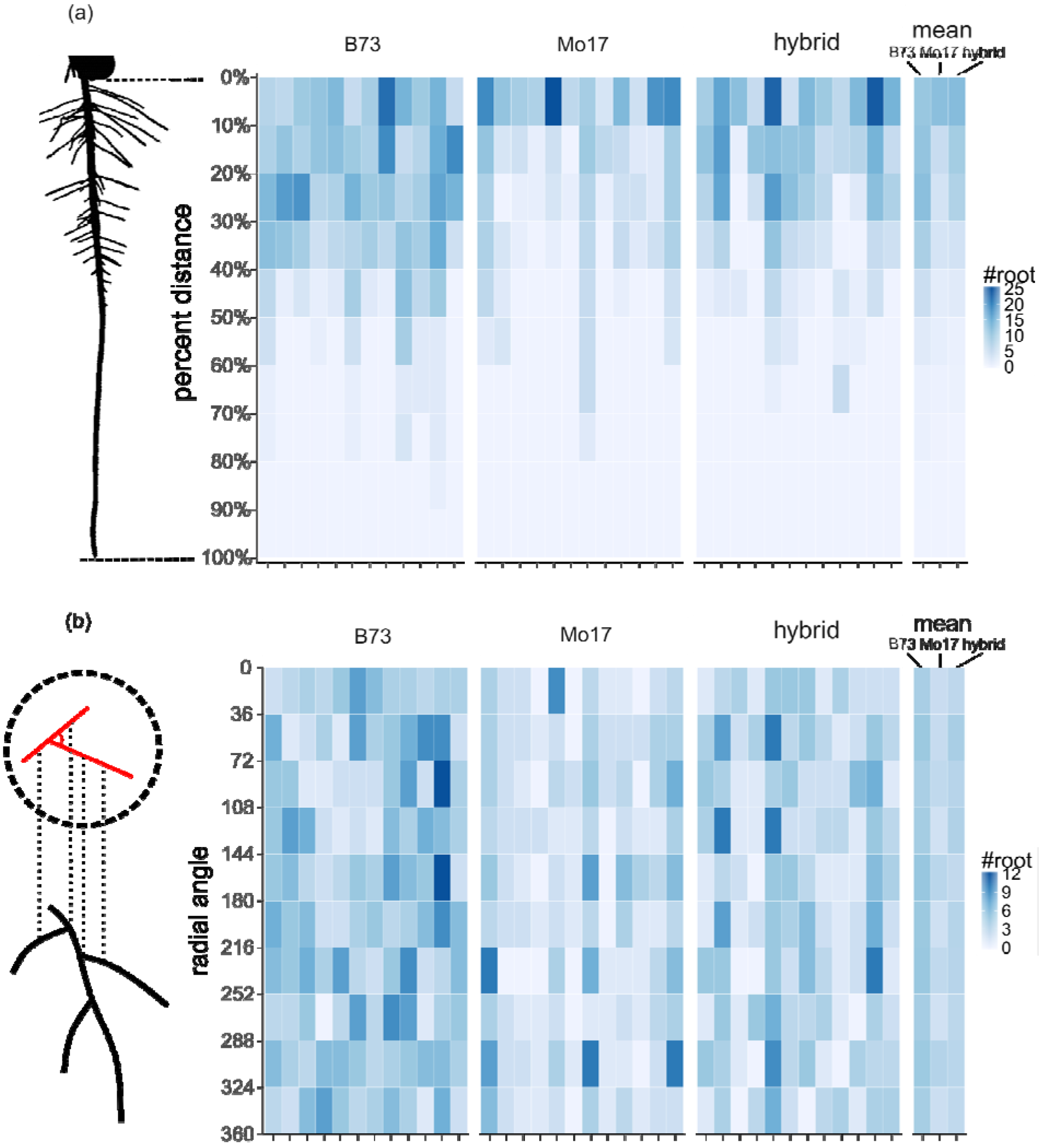
Developmental analysis along primary root revealing genotypic difference. (a) Distribution of the first-order lateral root number on the primary root. The percent distance refers to the distance between the branching fork to the primary root base, divided by the primary root length. (b) Distribution of the radial angles for lateral branching around the primary root. The radial angle is the angle formed by the root branch and the adjacent branch along the longitudinal axis. For each genotype, each column represents an individual seedling.

The radial position of new lateral roots is another developmental attribute that could influence 3D root system properties (Topp and Benfey 2011). Lateral roots develop from pericycle cells adjacent to vascular tissue. The well-studied 2D branching plane of *Arabidopsis thaliana* results from a simple diarch symmetry of vascular strands, but the polyarch symmetry of maize and other monocots promotes the radial emergence of lateral roots (60). To quantify if genotype-specific radial emergence patterns contributed to differences in global architecture, we leveraged our 3D data to compute radial angles for lateral branching around the primary root. Each lateral was compared to the adjacent branch along the longitudinal axis, and the relative angle between them was recorded in 10 bins, 36 degrees apart. The mean distributions of the radial angle had a near constant probability, suggesting that lateral roots overall emerged in all directions equally for B73, Mo17, and their hybrid (Figure 6b; mean values). However, the significant intra-plant variation across the experiment points to substantial heterogeneity along any given primary root, suggesting that radial angle is either stochastic or that micro-environmental conditioning may play a strong role.

## DISCUSSION

Here we quantified high-resolution spatiotemporal dynamics of maize seedling root systems in three dimensions as they rapidly grew in complexity. Mathematical modelling allowed us to evaluate the contributions of genotype by simple function-values that integrated growth trends along both real and developmental timescales. We demonstrate how differences in aggregate local growth dynamics and developmental characteristics of a single root can condition emergent global architectures, resulting in a highly branched, dense root system for B73 and a more open and extensive system for Mo17. Corresponding 3D X-ray-based analysis of field-excavated mature root crowns suggested these architectural properties were developmentally hardwired and environmentally stable.

When considered in the context of rhizoeconomic foraging strategy (61), the maize inbreds contrast for their carbon investments in new root production relative to maintenance of existing root growth - these differences have significant implications for resource acquisition. Previous studies have posited that the ideotype of fewer but longer lateral roots (Mo17) acquires mobile resources like nitrogen more efficiently, but the ideotype of finer but denser lateral roots (B73) acquires immobile resources like phosphorous more efficiently (15). Our findings extend the known P efficiency of B73 and inefficiency of Mo17 (62, 63) to include a basis in root growth dynamics. Work is underway to understand N relations. Through computational dissection of highly resolved spatiotemporal data, we have shown how genetic differences in complex developmental and local growth patterns measurable at the seedling level can be diagnostic of root architecture at maturity. Similar tradeoffs for thoroughness versus extent of soil exploration have been previously identified in rice seedling root systems, including genetic tradeoffs (64). Time-resolved phenotyping applied to mapping populations or association panels would resolve the genetic basis of complex root architecture dynamics, as it has for simpler root traits (33, 65). Such function-valued approaches are fundamentally superior than simple summary statistics to quantify growth and environmental interactions (66). Our model sensitivity analysis provides guidance on the imaging frequency that these studies could be effectively conducted.

With the information of local traits, the contribution of individual branches to the whole root system architecture can be studied. Despite a wide range of phenotypic variation within the root system of a single plant or among genetically identical plants, the overall patterns were strongly correlated with genotype. Considering the homogenous conditions of the gel growth system, we interpret this to mean that a significant amount of probabilistic behavior is inherent to root foraging, but these distributions may be genetically conditioned. Our multiscale approach can be extended to a wide variety of other research questions where local interactions, such as patchy nutrients or competing roots, can be studied in the context of their effects on global architecture. 3D imaging and computational analysis of root growth allows measurement of traits (e.g. radial angle) that are not measurable via 2D phenotyping approaches, especially with very complex root systems. It also allowed us to compare the 3D architectures of seedling and mature root crowns using the same algorithms. Our methods transfer to any 3D data, including the growing body of X-ray tomography (XRT) and magnetic resonance imaging (MRI) root studies (8, 47, 67, 68). Such high quality inputs should be especially useful to parameterize and constrain multiscale models that incorporate genotypic differences as probability distributions that reflect the inherent stochasticity in plant growth (19–22, 35, 57, 69–71). The continued improvement of simulation models that can account for and accurately predict root growth and root-environment interactions as a function of genotype will be critical for a realistic understanding of integrated plant biology and the enormous potential of root systems for the improvement of agriculture.

## METHODS

### Plant materials and growth conditions

Two maize inbred genotypes, B73 and Mo17, and their hybrid were used in this study. The growth medium was made following a modified Hoaglands solution (39). The seeds were sterilized with 35% hydrogen peroxide for 20 minutes and rinsed four times with RO water. After soaking in RO water for 8 hours at 29 °C in the dark, the seeds were sterilized again with 35% hydrogen peroxide for 10 minutes and rinsed four times with sterile water. The seeds were placed at 29 °C in the dark for incubation. After 2 days, seedlings were planted into glass growth cylinders sealed with saran-wrap. The cylinders were put on a dark shelf at ambient conditions overnight for acclimation before moving them into a growth chamber. The light intensity at the top of each jar was 700 µmol/m²/s. Humidity in the chamber was maintained at 50%, although the jars were sealed with saran wrap. Temperatures were set to 28 °C during the day and 24 °C at night, with a 16/8h day/night cycle.

### Imaging system

The imaging system was set up in the growth chamber. It consists of a digital camera (Stingray F-504C, Allied Vision Technologies), a turntable (LT360, LinearX), a near infrared LED light (SOBL, Smart Vision Lights), an optical correction tank, and a personal computer. The schematic details of a nearly identical imaging system can be seen in (38). The near infrared LED was used with a longpass camera filter to provide a high contrast silhouette during the day and to avoid affecting plant growth during night. It was turned on 10 seconds before each imaging course, and was turned off immediately after the rotational sequence was taken. For each imaging course, 180 images were taken at 2° increments. A custom program written in LabVIEW was used to control the image acquisition and set up time lapse imaging, which enabled the imaging system to take rotational image sequences at regular time points automatically. In this study, all the plants were imaged every 4 hours for a week, until day 11 after germination. For each plant, 41 or 42 image sets were captured.

### Inbred vs Hybrid Field Experiment

The same maize inbreds and hybrids that were used in the gel were included in a field experiment at the University of Missouri Genetics Farm in Columbia, MO planted on May 16, 2017. Six plants each were sampled from two rows after flowering (72), and washed root crowns were scanned on a North Star Imaging X5000 X-ray Computed Tomography system at the Danforth Plant Science Center. All the scans were performed at 70kV and 1700μA, collecting 1800 projections over 360 degrees of rotation. Scanning resolution was 111μm. The total scan time for each sample was 3 minutes. Projections were reconstructed into single 3D volumes using NSI efX-ct software, and each volume was exported as a 2D image stack for analysis. For the root segmentation, 2D image slices were thresholded using band thresholding to remove any soil that remained in the images. The two threshold values were determined by triangle algorithm and Otsu algorithm, respectively. Cleaned 3D root models were input into the RSA-GiA pipeline to calculate root trait values.

### Quantification of RSA traits

We used the RSA-GiA pipeline to generate the 3D reconstructions from 2D rotational images. The pipeline included three main steps: (1) cropping, to remove the above gel parts from the images; (2) thresholding, to convert the images to binary images, which roots are the foreground; (3) reconstruction, to build the 3D models based on visual hull algorithm (45). To analyze a time-series of 3D reconstructions, we used DynamicRoots. DynamicRoots is a software tool which is capable of computing structural and dynamic traits for growing roots (46). It aligns all the models in a time-series, decomposes the 3D root system into individual branches, and records the growth process. The primary output of DynamicRoots is a txt file that includes columns for different root traits at every observation time, and rows for data from every branch. To analyze root growth patterns, we developed R functions for computing global root traits, root growth rate, root growth direction, and root distribution from DynamicRoots-generated files (https://github.com/Topp-Roots-Lab/timeseries_analysis). Using the traits for each branch, the total volume, total length, and total number of branches were obtained for every time point. In order to avoid noise, branches shorter than 5 voxels at the last observation time were removed. The total root volume was the sum of the root volume of all individual roots. The total root length was the sum of the root length of all individual roots. The total number of branches was the sum of the numbers of individual roots. The growth rate was the difference of root traits between two subsequent time points. We defined the angle between branch and soil level as soil angle, and the angle between branch and its parent branch as branching angle.

### Data analysis

To compare the shape of root growth for different genotypes, we modeled the dynamic of global root traits. Linear, exponential, power law, monomolecular, three-parameter logistic, four-parameter logistic, and Gompertz models were tested. The basic functional forms for these models can be seen in Paine et al., 2012 (55). We parameterized the three-parameter logistic model in the following way to facilitate descriptions of growth curves:

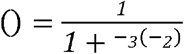

where *y* represents the global traits, i.e. total root volume, total root length, and total root number, *t* is the time after the start of the imaging course, α_1_ is the maximum growth capacity, α_2_ is the inflection time point, and α_3_ is the steepness.

The parameters for all models were estimated using linear (“lm” function) or non-linear least squares regression (“nls” function) in R. To select the best model, R squared (R^2^), root mean squared error (RMSE), standard error (S), and Akaike Information Criterion (AIC) of the regression were computed. t-test was used to compare the parameters among different genotypes.

## ACKNOWLEDGEMENTS

This material is based upon work supported by the National Science Foundation under Award numbers: IOS-1638507 and IIA-1355406.

We’d like to thank Dr. R. Kelly Dawe and members of the Topp Lab for critical reading and commentary on the manuscript.

## REFERENCES

1. Lynch J (1995) Root Architecture and Plant Productivity. Plant Physiol 109(1):7–13.

2. Doussan C, Pagès L, Vercambre G (1998) Modelling of the Hydraulic Architecture of Root Systems: An Integrated Approach to Water Absorption—Model Description. Ann Bot 81(2):213–223.

3. Lambers H, Shane MW, Cramer MD, Pearse SJ, Veneklaas EJ (2006) Root structure and functioning for efficient acquisition of phosphorus: Matching morphological and physiological traits. Ann Bot 98(4):693–713.

4. Hodge A, Berta G, Doussan C, Merchan F, Crespi M (2009) Plant root growth, architecture and function. Plant Soil 321(1-2):153–187.

5. Bao Y, et al. (2014) Plant roots use a patterning mechanism to position lateral root branches toward available water. Proc Natl Acad Sci U S A 111(25):9319–9324.

6. de Dorlodot S, et al. (2007) Root system architecture: opportunities and constraints for genetic improvement of crops. Trends Plant Sci 12(10):474–481.

7. Topp CN, Bray AL, Ellis NA, Liu Z (2016) How can we harness quantitative genetic variation in crop root systems for agricultural improvement? J Integr Plant Biol 58(3):213–225.

8. Morris EC, et al. (2017) Shaping 3D Root System Architecture. Curr Biol 27(17):R919–R930.

9. Muraya MM, et al. (2017) Genetic variation of growth dynamics in maize (Zea mays L.) revealed through automated non-invasive phenotyping. Plant J 89(2):366–380.

10. Zhang X, et al. (2017) High-Throughput Phenotyping and QTL Mapping Reveals the Genetic Architecture of Maize Plant Growth. Plant Physiol 173(3):1554–1564.

11. Weaver JE, Bruner WE (1926) Root development of field crops (McGraw-Hill Book Company New York and London).

12. Wachsman G, Sparks EE, Benfey PN (2015) Genes and networks regulating root anatomy and architecture. New Phytol 208(1):26–38.

13. Slovak R, Ogura T, Satbhai SB, Ristova D, Busch W (2016) Genetic control of root growth: from genes to networks. Ann Bot 117(1):9–24.

14. Gao Y, Lynch JP (2016) Reduced crown root number improves water acquisition under water deficit stress in maize (Zea mays L.). J Exp Bot 67(15):4545–4557.

15. Lynch JP (2013) Steep, cheap and deep: an ideotype to optimize water and N acquisition by maize root systems. Ann Bot 112(2):347–357.

16. Uga Y, et al. (2013) Control of root system architecture by DEEPER ROOTING 1 increases rice yield under drought conditions. Nat Genet 45(9):1097–1102.

17. Gamuyao R, et al. (2012) The protein kinase Pstol1 from traditional rice confers tolerance of phosphorus deficiency. Nature 488(7412):535–539.

18. Dupuy L, Gregory PJ, Bengough AG (2010) Root growth models: towards a new generation of continuous approaches. J Exp Bot 61(8):2131–2143.

19. Schnepf A, et al. (2018) CRootBox: a structural-functional modelling framework for root systems. Ann Bot. doi:10.1093/aob/mcx221.

20. Postma JA, et al. (2017) OpenSimRoot: widening the scope and application of root architectural models. New Phytol 215(3):1274–1286.

21. Kalogiros DI, et al. (2016) Analysis of root growth from a phenotyping data set using a density-based model. J Exp Bot 67(4):1045–1058.

22. Zhao J, et al. (2017) Root architecture simulation improves the inference from seedling root phenotyping towards mature root systems. J Exp Bot. doi:10.1093/jxb/erw494.

23. Spalding EP, Miller ND (2013) Image analysis is driving a renaissance in growth measurement. Curr Opin Plant Biol 16(1):100–104.

24. Downie HF, et al. (2015) Challenges and opportunities for quantifying roots and rhizosphere interactions through imaging and image analysis. Plant Cell Environ 38(7):1213–1232.

25. Johnson MG, Tingey DT, Phillips DL, Storm MJ (2001) Advancing fine root research with minirhizotrons. Environ Exp Bot 45(3):263–289.

26. Nagel KA, et al. (2012) GROWSCREEN-Rhizo is a novel phenotyping robot enabling simultaneous measurements of root and shoot growth for plants grown in soil-filled rhizotrons. Funct Plant Biol 39(11):891–904.

27. Iijima M, Oribe Y, Horibe Y, Kono Y (1998) Time Lapse Analysis of Root Elongation Rates of Rice and Sorghum During the Day and Night. Ann Bot 81(5):603–607.

28. Walter A, et al. (2002) Spatio-temporal dynamics of expansion growth in roots: automatic quantification of diurnal course and temperature response by digital image sequence processing. J Exp Bot 53(369):689–698.

29. Miller ND, Parks BM, Spalding EP (2007) Computer-vision analysis of seedling responses to light and gravity. Plant J 52(2):374–381.

30. French A, Ubeda-Tomás S, Holman TJ, Bennett MJ, Pridmore T(2009) High-throughput quantification of root growth using a novel image-analysis tool. Plant Physiol 150(4):1784–1795.

31. Yazdanbakhsh N, Fisahn J (2009) High throughput phenotyping of root growth dynamics, lateral root formation, root architecture and root hair development enabled by PlaRoM. Funct Plant Biol 36(11):938–946.

32. Lobet G, Pagès L, Draye X (2011) A novel image-analysis toolbox enabling quantitative analysis of root system architecture. Plant Physiol 157(1):29–39.

33. Moore CR, et al. (2013) High-throughput computer vision introduces the time axis to a quantitative trait map of a plant growth response. Genetics 195(3):1077–1086.

34. Rellán-Álvarez R, et al. (2015) GLO-Roots: an imaging platform enabling multidimensional characterization of soil-grown root systems. Elife 4. doi:10.7554/eLife.07597.

35. Passot S, et al. (2018) A New Phenotyping Pipeline Reveals Three Types of Lateral Roots and a Random Branching Pattern in Two Cereals. Plant Physiology. doi:10.1104/pp.17.01648.

36. Fang S, Yan X, Liao H (2009) 3D reconstruction and dynamic modeling of root architecture in situ and its application to crop phosphorus research. Plant J 60(6):1096–1108.

37. Iyer-Pascuzzi AS, et al. (2010) Imaging and analysis platform for automatic phenotyping and trait ranking of plant root systems. Plant Physiol 152(3):1148–1157.

38. Clark RT, et al. (2011) Three-dimensional root phenotyping with a novel imaging and software platform. Plant Physiol 156(2):455–465.

39. Zurek PR, Topp CN, Benfey PN (2015) Quantitative trait locus mapping reveals regions of the maize genome controlling root system architecture. Plant Physiol 167(4):1487–1496.

40. Mooney SJ, Pridmore TP, Helliwell J, Bennett MJ (2012) Developing X-ray Computed Tomography to non-invasively image 3-D root systems architecture in soil. Plant Soil 352(1-2):1–22.

41. Mairhofer S, et al. (2012) RooTrak: automated recovery of three-dimensional plant root architecture in soil from x-ray microcomputed tomography images using visual tracking. Plant Physiol 158(2):561–569.

42. Borisjuk L, Rolletschek H, Neuberger T (2012) Surveying the plant’s world by magnetic resonance imaging. Plant J 70(1):129–146.

43. Metzner R, et al. (2015) Direct comparison of MRI and X-ray CT technologies for 3D imaging of root systems in soil: potential and challenges for root trait quantification. Plant Methods 11:17.

44. Jahnke S, et al. (2009) Combined MRI–PET dissects dynamic changes in plant structures and functions. Plant J 59(4):634–644.

45. Topp CN, et al. (2013) 3D phenotyping and quantitative trait locus mapping identify core regions of the rice genome controlling root architecture. Proc Natl Acad Sci U S A 110(18):E1695–704.

46. Symonova O, Topp CN, Edelsbrunner H (2015) DynamicRoots: A Software Platform for the Reconstruction and Analysis of Growing Plant Roots. PLoS One 10(6):e0127657.

47. van Dusschoten D, et al. (2016) Quantitative 3D Analysis of Plant Roots Growing in Soil Using Magnetic Resonance Imaging. Plant Physiol 170(3):1176–1188.

48. Hufnagel B, et al. (2014) Duplicate and conquer: multiple homologs of PHOSPHORUS-STARVATION TOLERANCE1 enhance phosphorus acquisition and sorghum performance on low-phosphorus soils. Plant Physiol 166(2):659–677.

49. Wissuwa M, Kretzschmar T, Rose TJ (2016) From promise to application: root traits for enhanced nutrient capture in rice breeding. J Exp Bot 67(12):3605–3615.

50. Poorter H, et al. (2016) Pampered inside, pestered outside? Differences and similarities between plants growing in controlled conditions and in the field. New Phytol 212(4):838–855.

51. Hund A, Reimer R, Stamp P, Walter A (2012) Can we improve heterosis for root growth of maize by selecting parental inbred lines with different temperature behaviour? Philos Trans R Soc Lond B Biol Sci 367(1595):1580–1588.

52. Walter A, Silk WK, Schurr U (2009) Environmental effects on spatial and temporal patterns of leaf and root growth. Annu Rev Plant Biol 60:279–304.

53. Bray AL, Topp CN (2018) The Quantitative Genetic Control of Root Architecture in Maize. Plant Cell Physiol. doi:10.1093/pcp/pcy141.

54. Barlow PW, Brain P, Parker JS (1991) Cellular Growth in Roots of a Gibberellin-Deficient Mutant of Tomato (Lycopersicon esculentum Mill.) and its Wild-type. J Exp Bot 42(3):339–351.

55. Paine CET, et al. (2012) How to fit nonlinear plant growth models and calculate growth rates: an update for ecologists. Methods Ecol Evol 3(2):245–256.

56. Morris AK, Silk WK (1992) USE OF A FLEXIBLE LOGISTIC FUNCTION TO DESCRIBE AXIAL GROWTH OF PLANTS. Bulletin of Mathematical Biology 54(6):1069–1081.

57. Araya T, Kubo T, von Wirén N, Takahashi H (2016) Statistical modeling of nitrogen-dependent modulation of root system architecture in Arabidopsis thaliana. J Integr Plant Biol 58(3):254–265.

58. Postma JA, Dathe A, Lynch JP (2014) The optimal lateral root branching density for maize depends on nitrogen and phosphorus availability. Plant Physiol 166(2):590–602.

59. Petricka JJ, Winter CM, Benfey PN (2012) Control of Arabidopsis root development. Annu Rev Plant Biol 63:563–590.

60. Smith S, De Smet I (2012) Root system architecture: insights from Arabidopsis and cereal crops. Philos Trans R Soc Lond B Biol Sci 367(1595):1441–1452.

61. Lynch JP, Ho MD, Phosphorus L (2005) Rhizoeconomics: Carbon costs of phosphorus acquisition. Plant Soil 269(1-2):45–56.

62. Kaeppler SM, et al. (2000) Variation among Maize Inbred Lines and Detection of Quantitative Trait Loci for Growth at Low Phosphorus and Responsiveness to Arbuscular Mycorrhizal Fungi Work supported by USDA-Hatch, University of Wisconsin University-Industry Relations, and Cargill Fertilizer. Crop Sci 40(2):358–364.

63. Zhu J, Kaeppler SM, Lynch JP (2005) Mapping of QTLs for lateral root branching and length in maize (Zea mays L.) under differential phosphorus supply. Theor Appl Genet 111(4):688–695.

64. Topp CN, et al. (2013) 3D phenotyping and quantitative trait locus mapping identify core regions of the rice genome controlling root architecture. Proc Natl Acad Sci U S A 110(18):E1695–E1704.

65. Kwak I-Y, Moore CR, Spalding EP, Broman KW (2014) A simple regression-based method to map quantitative trait loci underlying function-valued phenotypes. Genetics 197(4):1409–1416.

66. Stinchcombe JR, Function-valued Traits Working Group, Kirkpatrick M (2012) Genetics and evolution of function-valued traits: understanding environmentally responsive phenotypes. Trends Ecol Evol 27(11):637–647.

67. Mooney SJ, Pridmore TP, Helliwell J, Bennett MJ (2012) Developing X-ray Computed Tomography to non-invasively image 3-D root systems architecture in soil. Plant Soil 352(1-2):1–22.

68. Ahmed S, et al. (2016) Imaging the interaction of roots and phosphate fertiliser granules using 4D X-ray tomography. Plant Soil 401(1):125–134.

69. Lynch JP, Nielsen KL, Davis RD, Jablokow AG (1997) SimRoot: Modelling and visualization of root systems. Plant Soil 188(1):139–151.

70. Dupuy L, Gregory PJ, Bengough AG (2010) Root growth models: towards a new generation of continuous approaches. J Exp Bot 61(8):2131–2143.

71. Bodner G, et al. (2013) A statistical approach to root system classification. Front Plant Sci 4:292.

72. Trachsel S, Kaeppler SM, Brown KM, Lynch JP (2011) Shovelomics: high throughput phenotyping of maize (Zea mays L.) root architecture in the field. Plant Soil 341(1):75–87.

